# An integrative model of plant gravitropism linking statoliths position and auxin transport

**DOI:** 10.1101/2021.01.01.425032

**Authors:** Nicolas Levernier, Olivier Pouliquen, Yoël Forterre

**Affiliations:** Aix Marseille Univ, CNRS, IUSTI, Marseille, France

**Keywords:** Plant tropism, Gravity sensing, Auxin signaling, PIN trafficking, Modeling

## Abstract

Gravity is a major cue for the proper growth and development of plants. The response of plants to gravity implies starch-filled plastids, the statoliths, which sediments at the bottom of the gravisensing cells, the statocytes. Statoliths are assumed to modify the transport of the growth hormone, auxin, by acting on specific auxin transporters, PIN proteins. However, the complete gravitropic signaling pathway from the intracellular signal associated to statoliths to the plant bending is still not well understood. In this article, we build on recent experimental results showing that statoliths do not act as gravitational force sensor, but as position sensor, to develop a bottom-up theory of plant gravitropism. The main hypothesis of the model is that the presence of statoliths modifies PIN trafficking close to the cell membrane. This basic assumption, coupled with auxin transport and growth in an idealized tissue made of a one-dimensional array of cells, recovers several major features of the gravitropic response of plants. First, the model provides a new interpretation for the response of a plant to a steady stimulus, the so-called sine-law of plant gravitropism. Second, it predicts the existence of a gravity-independent memory process as observed recently in experiments studying the response to transient stimulus. The model suggests that the timescale of this process is associated to PIN turnover, calling for new experimental studies.

## 1 INTRODUCTION

The detection of gravity by plants and the resulting growth response (gravitropism) offer a fascinating illustration of a multi-scale perception mechanism in living organisms (Fig. 1) (Moulia and Fournier, 2009; Morita, 2010; Toyota and Gilroy, 2013). It originates in specific cells, called statocytes, where tiny starch-accumulating amyloplasts acting as statoliths sediment under gravity at the bottom of the cells (Fig. 1C). When the plant is inclined, the repositioning of statoliths under gravity induces a relocalisation of auxin transporters (PIN proteins) at the membrane of statocytes, which generates a lateral transport of auxin toward the lower side of the shoot or the root (Cholodni–Went hypothesis) (Fig. 1B). In turn, this asymmetry in auxin concentration induces a differential growth across the plant organ, and thus its bending toward the gravity vector (Fig. 1A). Since the pioneering works of the Darwins and Sachs on plant tropisms (Darwin and Darwin, 1880; Sachs, 1887), progress has been made on every step of this gravitropic signaling pathway. Yet, basic questions remain unanswered. In particular, it is still not clear how the first physical signal generated by the sedimentation of statoliths is converted into biochemical signals downstream, to eventually produce the growth response at the plant scale (Nakamura et al., 2019).

**Figure 1.**
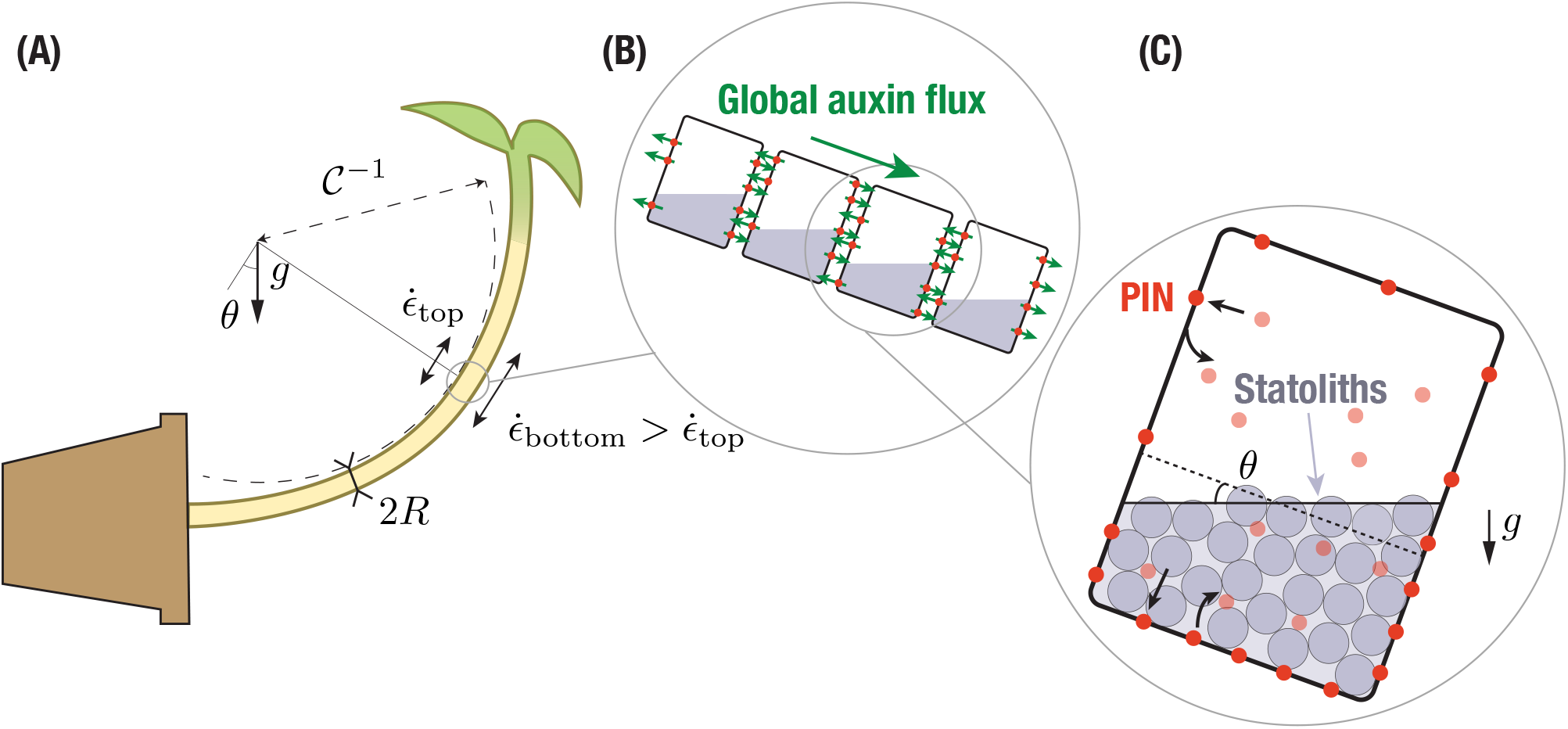
Multiscale description of gravitropism. At the macroscopic scale **(A)**, the response to gravity of a shoot or a stem is achieved by differential growth across the organ, which induces a curvature of the organ. At the tissue scale **(B)**, differential growth results from a net flux of the auxin across the width (large green arrows), owing to the asymmetric distribution of auxin transporters (PINs, red circles). The local auxin fluxes are shown by the small green arrows. At the cell scale **(C)**, PIN asymmetry results from the asymmetric distribution of the statoliths position after sedimentation under gravity, which modifies PIN trafficking close to the cell membrane.

Recently, insights into the sensing mechanism and the transduction pathway have been obtained from experiments both at the macroscopic and microscopic levels. First, the gravitropic response to permanent stimuli (inclination of the plant), the so-called sine-law of gravitropism (Sachs, 1887; Larsen, 1969; Iino et al., 1996; Galland, 2002; Dumais, 2013), was found to depend on the inclination but, surprisingly, not on the intensity of gravity (Chauvet et al., 2016). Hence, statocytes behave like inclination sensors not force sensors as previously believed. An important consequence is that the initial gravity stimulus for gravitropism should be the position of the statoliths within statocytes (Pouliquen et al., 2017). This position-sensor hypothesis gained a mechanistic support from the direct observation of statoliths motion under gravity stimulation (Bérut et al., 2018). Unlike a pile of macroscopic grains like sand, statoliths were found to move and flow at any inclination. This liquid-like behavior comes from the random agitation of the statoliths, whose origin is not thermal but arises from the interaction of statoliths with the acto-myosin cytoskeleton inside the cell (Sack et al., 1986; Saito et al., 2005; Nakamura et al., 2011). A second insight came from dose-response like experiments on wheat coleoptiles, in which the gravity stimulus was applied during a transient period only (Chauvet et al., 2019). When the shoots were inclined for short period of time, the gravitropic response was found to deviate from the steady response and decay. The transition occurred for a time *τ*_memory_ ~ 15 min, which was independent of gravity and much larger than the statoliths sedimentation time. This observation suggested the existence of a memory-integration process in the gravitropic signaling pathway, independent of the statoliths dynamics, which integrates the initial signal induced by statoliths displacement.

To account for these observations (position-sensor hypothesis, memory time independent of *g*), Chauvet et al. (2019) built a mathematical model of gravitropism in which the gravitropic signal controlling the differential growth was linked to the statoliths position by an integrative process of timescale *τ*_memory_ (a similar approach was used in Meroz et al. (2019)). Once coupled to the statoliths dynamics and the tropic growth motion, the model was able to reproduce the transient gravitropic response observed experimentally. However, Chauvet et al. (2019)’s model was built on two *ad-hoc* postulates. First, it assumed that the relation between the gravitropic signal and the statoliths position is known and given by the sine-law. Second, it postulated the existence of the integrative process and time scale *τ*_memory_, without explaining its origin. The spatio-temporal dynamics of the molecular processes acting between the statoliths and the growth response, such as the dynamics of PIN proteins and auxin transport, was not described.

The objective of this paper is to fill this gap by building an integrative model of plant gravitropism that bridges the different scales of the process: (i) the initial intracellular gravitropic signal encoded in the statoliths position, (ii) PINs dynamics at the cellular level, (ii) auxin transport at the tissue level and, finally, (iv) differential growth and curvature at the plant organ scale. Previous models of plant gravitropism mainly focused either on the macroscopic scale, describing how the complex spatio-temporal evolution of the organ shape results from the interplay between differential growth and the slender geometry of the organ (Bastien et al., 2013, 2014; Chelakkot and Mahadevan, 2017), or on the tissue level, modeling growth mechanics (Dyson et al., 2014) or auxin transport (Band et al., 2012; Fendrych et al., 2018; Retzer et al., 2019) in realistic 1D or 3D tissue geometries. In these latter models, the distribution of PINs in response to plant inclination was prescribed and not linked to the intracellular dynamics of the statoliths. This is precisely the goal of our study. Building on the recent position-sensor hypothesis, we propose a simple but generic model of interaction between statoliths and PINs trafficking at the cell membrane, that we couple with the classical equations of auxin transport and tissue growth. We then study the gravitropic response predicted by the model for steady and unsteady gravity stimuli, comparing the results with the experiments of Chauvet et al. (2016) and Chauvet et al. (2019).

## 2 MATERIAL AND METHODS

### 2.1 Link between gravitropic curvature, differential growth and auxin concentration gradient

At the plant scale, the gravitropic response is characterized by the curvature of the organ resulting from differential growth, which itself results from auxin gradients (Cholodny–Went hypothesis). The first step of the model is thus to relate those three quantities. For a slender organ like a shoot or a stem, the rate of change of the local curvature 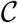 is related to differential growth through the following kinematic relationship: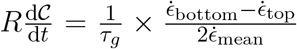, where *R* is the radius of the organ, 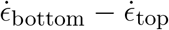 is the difference of growth rate between both sides, 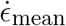 is the mean growth rate and 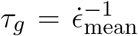 is the growth timescale (Silk, 1984; Moulia and Fournier, 2009; Bastien et al., 2013) (Fig. 1). The growth rate of plant cells is known to be controlled by auxin, the so-called growth hormone. Auxin stimulates cell elongation by loosening cell walls. To the best of our knowledge, the link between the local auxin concentration in walls and the local growth rate of cells has not been robustly determined and only the response of the whole tissue to an external addition of auxin has been investigated. It is however often assumed that growth is mainly controlled by the auxin concentration in the vicinity of the ‘skin’ of the organ, as epidermal tissues are stiffer than inner tissues (Kutschera and Niklas, 2007; Dyson et al., 2014). For the sake of simplicity, we will here assume that the local growth rate is simply proportional to the local auxin concentration *c*, 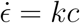 (Galston and Hand, 1949; Hopkins and Hüner, 2009), such that:

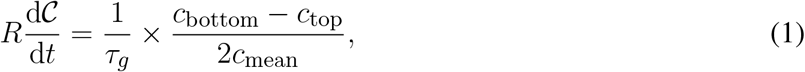

where 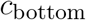 and 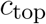 are the auxin concentrations on both sides of the organ and 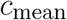 the mean auxin concentration. Under this assumption, the dimensionless gravitropic response deduced from the curvature dynamics, 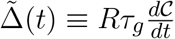, is equal to the relative auxin gradient across the organ, 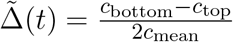. The goal of the model is to predict how this auxin gradient establishes when the plant is tilted.

### 2.2 Auxin transport

Auxin transport plays a key role in shaping plants development and, as such, has been the topic of extensive research over the past decades. Auxin transport is based on two distinct mechanisms (Goldsmith, 1977; Hopkins and Hüner, 2009; Runions et al., 2014). On the one hand, auxin in cell walls (mostly in a protonated form) enters the neighboring cell passively, or thanks to Aux/Lax influx carriers that are evenly distributed throughout the membrane. On the other hand, auxin inside cells (mostly in an anionic form) can only exit thanks to active auxin efflux carriers, such as PIN proteins (Krecek et al., 2009) or ABCB transporters (Zažímalová et al., 2010). While ABCB are evenly distributed throughout the membrane, PIN proteins are usually polarized and can be redistributed in response to external stimuli such as gravity (in particular PIN3, which is known to be implied in gravitropic response, see Friml et al. (2002)). Hence, an asymmetric distribution of PIN carriers on each side of the cell can generate an active transport of auxin from one cell to the other, resulting in a stable auxin gradient.

To model this situation, we provide a simplified description of auxin transport in which the different forms of auxin (proton-associated or not) are not taken into account. The tissue across the shoot or stem (width 2*R*) is modeled as a one-dimensional array of *N* cells of width *W*, separated by a cell wall of width *w* (Fig. 1 b and Fig. 2). We denote *c*_*n*_ the auxin concentration inside the *n*-th cell and *C*_*n*_ the auxin concentration inside the *n*-th wall, which are both assumed uniform (the equilibrium time of auxin in each compartment is very fast, about 0.1 s in the cell wall and few s inside the cell taking typical values of auxin diffusion coefficients, see Kramer et al. (2007)). We also neglect auxin dilution due to cell growth and assume that auxin is neither degraded nor created, as the degradation time of auxin (of the order of the day, see Grieneisen et al. (2012)) is much longer than the minutes to hours timescales we are interested in. The efflux current of auxin (number of auxin molecules per unit time and unit surface) from the *n*-th cell to the left wall (resp. right wall) is given by 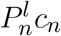 (resp. 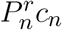), where 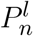 (resp. 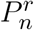) is the permeability of the left (resp. right) membrane (unit m/s). Conversely, the influx current of auxin from the *n*-th wall to both adjacent cells is *P*^in^*C*_*n*_, where the influx permeability *P*^in^ is assumed uniform for all cells (see Fig. 2). The time-evolution of the concentration is then:

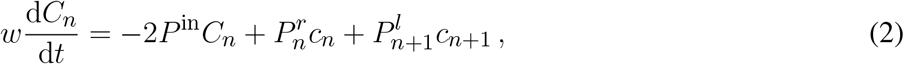

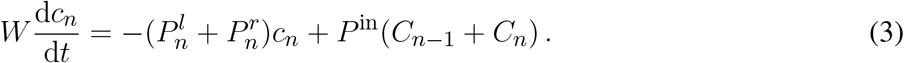

**Figure 2.**
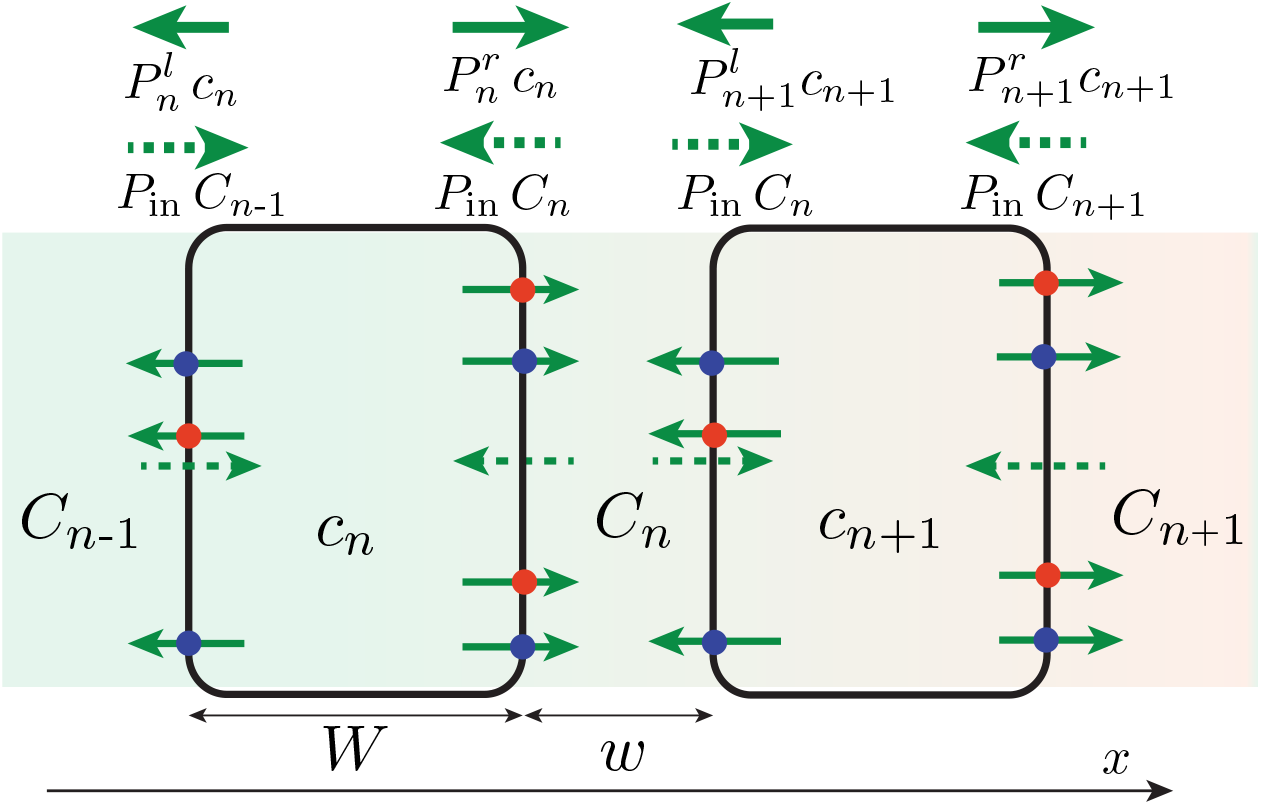
One-dimensional, discrete model of auxin transport across the tissue (in reality *w* ≪ *W*). Efflux of auxin (solid green arrow) occurs through efflux carriers (PIN: red circle, ABCB: blue circle), whose distribution (and thus permeabilities 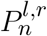) can be different on the right and left membrane of the cell. By contrast, influx of auxin (green dotted arrow) occurs with a symmetrical permeability *P*^in^ on both side of the cell. An asymmetry of efflux permeabilities *P*^*l*^ ≠ *P*^*r*^ can generate a net flux of auxin across the tissue, yielding an auxin concentration gradient (background color gradient).

The cell wall size being much smaller than the cell width (*w* ≪ *W*), the auxin concentration in the cell wall can be assumed quasi-steady, 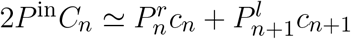, yielding:

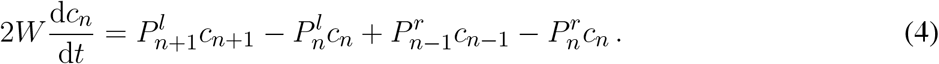

In the following, we assume that the distribution of auxin efflux carriers is the same in each cell, so that *P*^*l*^ and *P*^*r*^ are independent of *n*. This is the case of shoot coleoptiles where all cells in the growing region are similar and contain statoliths, but not the case of stems like the inflorescence of *Arabidopsis*, where statoliths are only present on an external ring in the endodermal cells (the modification of the equation in this case of inhomogeneous tissue is given in Appendix A). We also assume that auxin gradients occur over a length scale much larger than the cell size. In the continuum limit [*c*_*n*_(*t*) → *c*(*x, t*)], equation (4) for auxin transport then reduces to an advection-diffusion equation given by:

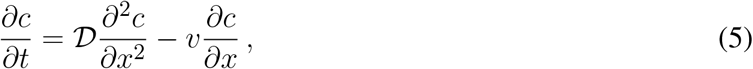

where 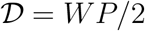 is the coefficient of diffusion and *v* = *δP*/2 the advection speed, with *P* = (*P*^*l*^ + *P*^*r*^)/2 and *δP* = *P*^*r*^ − *P*^*l*^.

The advective part of equation (5), which is responsible for auxin transport from one side to the other and thus to the differentiated growth and the curvature of the organ, is entirely controlled by the asymmetry of efflux permeabilities *δP*. Since ABCB carriers are evenly distributed, this asymmetry comes solely from the asymmetry of PINs distribution between the right and left side of the cells:

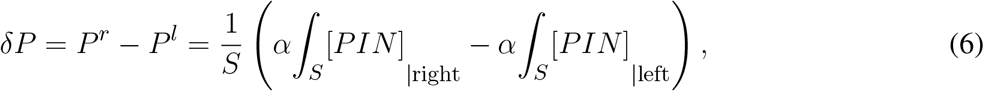

where *α* is the conductance of a single PIN carrier (unit m^3^/s), *S* the lateral surface of the cells and [*PIN*] the surface concentration of PINs attached to the membrane. In the following, we assume that the efflux permeability due to ABCB carriers is much larger than the efflux permeability due to PINs, such that *P* = (*P*^*l*^ + *P*^*r*^)/2 ≃ (1/*S*)*β* ∫ [*ABCB*], where *β* is the conductance of a single ABCB carrier, is independent of PINs concentration. This enables us to take a constant coefficient of diffusion 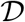 in the model, which simplify the results without affecting much the conclusions.

### 2.3 Coupling PIN dynamics to statoliths position: biased efflux at cell scale

The previous section relates auxin transport to the asymmetry of PINs distribution at the cellular level. We now model how this asymmetry emerges when the plant is tilted under gravity. Recently, it has been demonstrated that the relevant gravitropic stimulus for graviperception is the statoliths position within the statocytes (position-sensor hypothesis), and not their weight as previously believed (Chauvet et al., 2016; Pouliquen et al., 2017). Statoliths have also been identified as key actors in the relocalisation of PIN-proteins in response to change of gravity direction (Nakamura et al., 2019). Yet, how statoliths position is detected and read to modify PIN polarity remains largely unknown. PINs trafficking involves synthesis in the endoplasmic reticulum, degradation in the vacuole and recycling (Kleine-Vehn and Friml, 2008). Recycling is achieved by endocytosis, i.e., the deallocation of PIN proteins formerly attached to the cell membrane toward the cytoplasm inside a vesicle, or by exocytosis, i.e., the reallocation of the vesicle-carried PINs from the cytoplasm back to the cell membrane.

Following the position-sensor hypothesis, we assume that the presence of statoliths, either through direct steric constraints or through indirect molecular signaling, modify the trafficking of PIN proteins, so that PINs tend to enter the cell membrane preferentially on places where statoliths are in contact with it. This mechanism is formalized as follows. The endocytosis rate of PINs, d[*PIN*]_*i*_/d*t*|_endo_ = −*k*_off,*i*_[*PIN*]_*i*_ where [*PIN*]_*i*_ is the surface concentration of PIN attached to the membrane, is assumed to depend on the presence of statoliths, with *i* = 0 if no statoliths are present and *i* = 1 if they are (see Fig. 3). Similarly, the rate of exocytosis is written as d[*PIN*]_*i*_/d*t*|_exo_ = +*k*_on,*i*_[*PIN*]_vol_, where [*PIN*]_vol_ is the volumic concentration of PINs molecules inside the cell of volume *WS*. Two cases will be distinguished in the model, depending on whether PINs can attach to any side of the cell (“apical/basal/lateral binding”) or only on lateral sides (“lateral binding”) (see Fig. 4A). Assuming that the total number *N*_tot_ of PINs is conserved during gravistimulation (Kleine-Vehn et al., 2010) leads to the following set of equations for the PIN concentration attached to the membrane, [*PIN*]_*i*=0,1_:

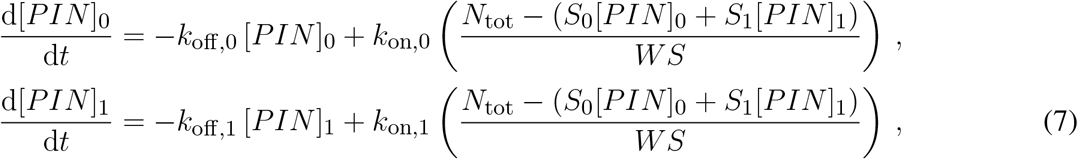

where *S*_1_ (resp. *S*_0_) denotes the total surface in contact (resp. not in contact) with statoliths in case of apical/basal/lateral binding, or only the lateral surfaces in contact (resp. not in contact) with statoliths in the lateral binding case.

**Figure 3.**
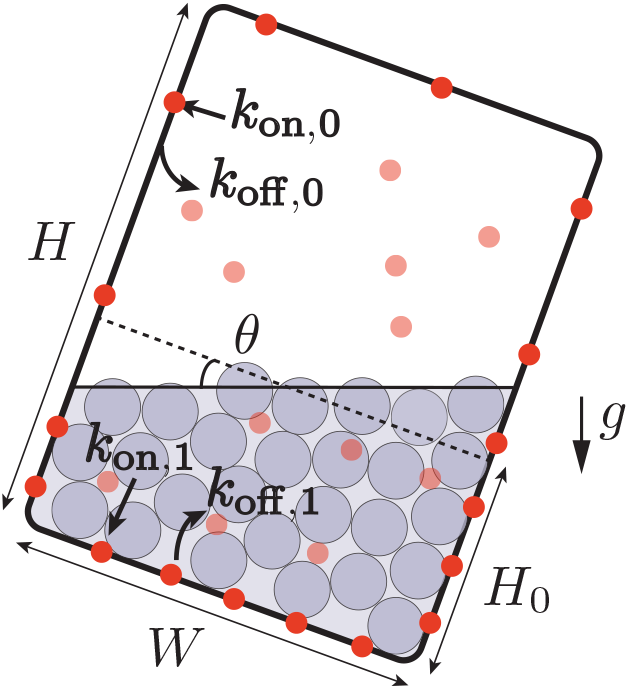
Interaction between PIN trafficking and statoliths position. The rate of reallocation *k*_on_ and deallocation *k*_off_ of PINs (bold red circles: PINs attached to the cell membrane, light red circles: PINs in bulk) depends on the presence of statoliths (grey).When the cell is tilted, the asymmetric distribution of the position of the statoliths induces a bias in the distribution of the PINs attached to the membrane.

**Figure 4.**
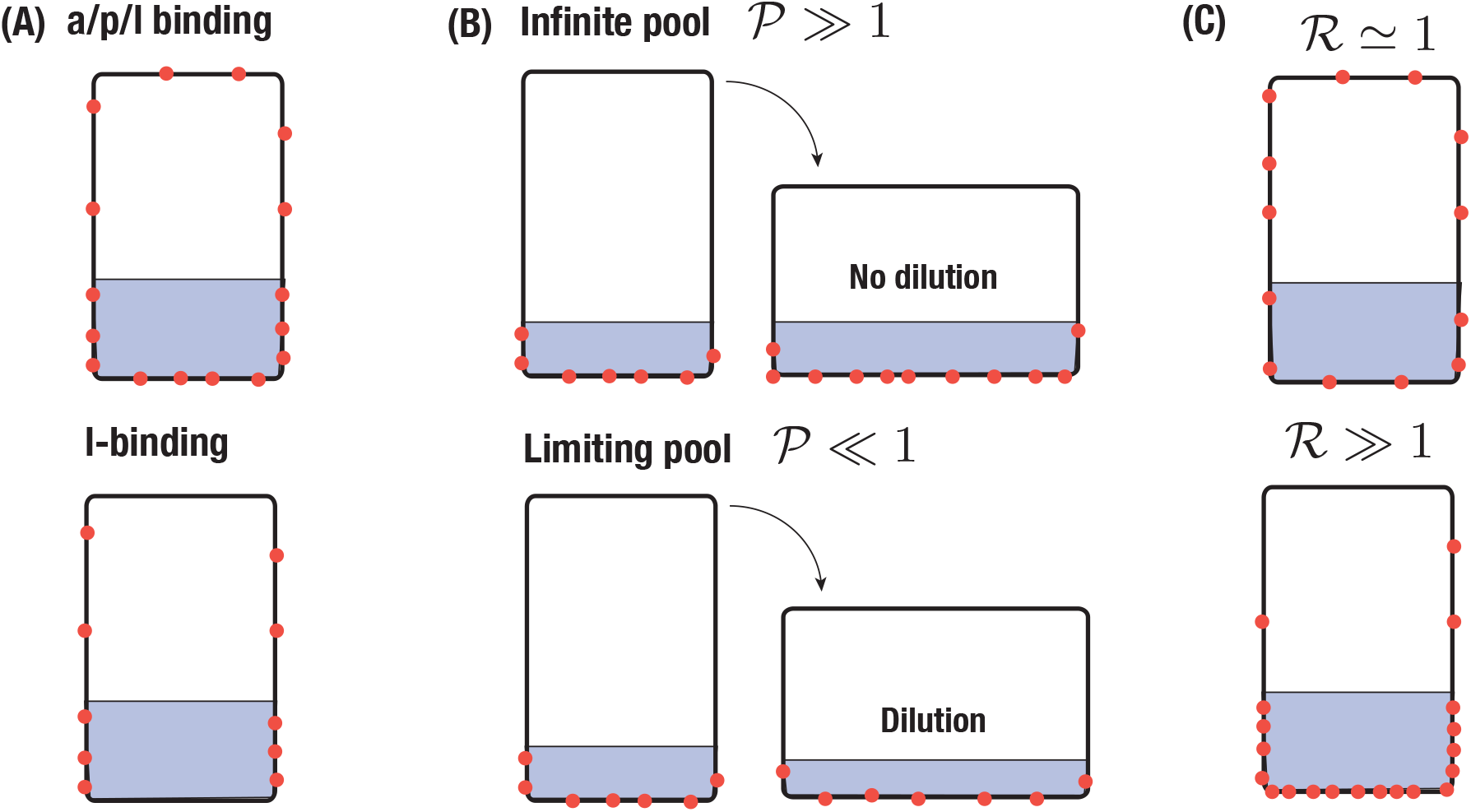
Sketch of different scenario of PIN-binding. PINs are represented in red and the region with statoliths in grey. **(A)** Apical/basal/lateral-binding vs lateral-binding. **(B)** Infinite pool versus limiting pool. In the first case, the surface density of PINs is conserved whereas in the second one, the total number of PINs is conserved. **(C)** Low sensitivity of PIN to statoliths 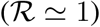 or high sensitivity 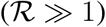.

The form of equation (7) shows that two regimes can be distinguished, depending on whether *W* is large or small compared to *k*_on_*/k*_off_. The first regime, called ‘infinite-pool’ regime in the following, corresponds to the case where *Wk*_off_ */k*_on_ is so large that binding is only limited by *k*_off_. The second regime, called ‘limiting-pool’ regime, corresponds to the opposite situation where the total number of attached carriers is almost equal to *N*_tot_: *S*_0_[*PIN*]_0_ + *S*_1_[*PIN*]_1_ ≃ *N*_tot_ (see Fig. 4B). Finally, we note that in writing equation (7), we have neglected the diffusion of PINs inside the membrane. This is justified since, over the time scales we are interesting in (running from minutes to one hour), PINs diffuse only over a distance of about few micrometers, which is much smaller than the cell size (taking 0.1*μm*^2^ min for the diffusion coefficient of a PIN, see Kleine-Vehn et al. (2011)).

Equations (1, 5, 7) represent a complete model describing the time-evolution of the gravitropic response once the statoliths is known, from the PIN localization to auxin transport and bending of the plant. In the following, we study the predictions of the model for a steady and transient stimulus. Table 1 and 2 summarizes all the physical quantities and dimensionless parameters defined from the model.

**Table 1.** List of dimensional quantities in the model with their definition and unit.

| Physical quantities | Definition | Unit |
| --- | --- | --- |
| $R$ | radius of the organ | (m) |
| $\mathcal{C}$ | bending curvature of the organ | (m<sup>-1</sup>) |
| $W$ | cell width | (m) |
| $w$ | cell wall width | (m) |
| $H$ | cell height | (m) |
| $H_0$ | statolith pile height before inclination | (m) |
| $S$ | cell lateral surface | (m<sup>2</sup>) |
| $S_{0,1}$ | total surface of the cell not (0) or (1) in contact with statoliths | (m<sup>2</sup>) |
| $S_0^{l,r}$ | surface of the left ($l$) or right ($r$) side of the cell not in contact with statoliths | (m<sup>2</sup>) |
| $S_1^{l,r}$ | surface of the left ($l$) or right ($r$) side of the cell in contact with statoliths | (m<sup>2</sup>) |
| $\dot{\epsilon}$ | growth rate | (s<sup>-1</sup>) |
| $\tau_g$ | $\tau_g = \dot{\epsilon}_{\text{mean}}^{-1}$ growth timescale | (s) |
| $c$ | auxin concentration inside the cell | (mol m<sup>-3</sup>) |
| $C$ | auxin concentration in the cell wall | (mol m<sup>-3</sup>) |
| $P^{l,r}$ | auxin efflux permeability of the left ($l$) or right ($r$) side of the cell | (m<sup>-1</sup>) |
| $P^{\text{in}}$ | auxin influx permeability | (m<sup>-1</sup>) |
| $\alpha$ | conductance of a single PIN carrier | (m<sup>3</sup> s<sup>-1</sup>) |
| $\beta$ | conductance of a single ABCB carrier | (m<sup>3</sup> s<sup>-1</sup>) |
| $P$ | $P = \frac{P^l + P^r}{2} = \frac{1}{S} \beta \int [ABCB]$ | (m<sup>-1</sup>) |
| $\delta P$ | $\delta P = P^r - P^l = \frac{1}{S} \left( \alpha \int_S [PIN]_{\text{right}} - \alpha \int_S [PIN]_{\text{left}} \right)$ | (m<sup>-1</sup>) |
| $\mathcal{D}$ | auxin coefficient of diffusion $\mathcal{D} = \frac{WP}{2}$ | (m<sup>2</sup> s<sup>-1</sup>) |
| $v$ | auxin advection speed $v = \frac{\delta P}{2}$ | (m<sup>-1</sup>) |
| $k_{\text{off},0,1}$ | endocytosis rate when statoliths are (1) or are not (0) in contact with the membrane | (s<sup>-1</sup>) |
| $k_{\text{on},0,1}$ | exocytosis speed when statoliths are (1) or are not (0) in contact with the membrane | (m s<sup>-1</sup>) |
| $\tau_{\text{aux}}$ | timescale for auxin transport across the tissue $\tau_{\text{aux}} = \frac{1}{\pi^2 + (\text{Pe}^2/4)} \times \frac{(2R)^2}{\mathcal{D}}$ | (s) |
| $\tau_{\text{PIN}}$ | timescale for PIN turnover $\tau_{\text{PIN}} = \frac{k_{\text{on},0}S_0 + k_{\text{on},1}S_1}{k_{\text{off},0}k_{\text{on},1}S_1 + k_{\text{off},1}k_{\text{on},0}S_0}$ | (s) |

**Table 2.** List of the dimensionless parameters used in the model with their definition and meaning.

| Dimensionless parameters | Definition | Meaning |
| --- | --- | --- |
| $\theta$ | | Inclination of the plant |
| $N$ | | Number of cells across the tissue |
| $N_{\text{tot}}$ | | Total number of PIN carriers per cell |
| $\tilde{\Delta}$ | $\tilde{\Delta} = \frac{c_{\text{bottom}} - c_{\text{top}}}{2c_{\text{mean}}}$ | Gravitropic response |
| $\text{Pe}$ | $\text{Pe} = \frac{2Rv}{\mathcal{D}}$ | Peclet number comparing auxin advection to diffusion |
| $\mathcal{A}$ | $\mathcal{A} = \frac{\alpha N_{\text{tot}} N}{\beta \int [\text{ABCB}]}$ | Effect of PIN conductance vs ABCB conductance on auxin transport |
| $\mathcal{R}$ | $\mathcal{R} = \frac{[\text{PIN}]_1^{\text{steady}}}{[\text{PIN}]_0^{\text{steady}}} = \frac{k_{\text{on},1} k_{\text{off},0}}{k_{\text{on},0} k_{\text{off},1}}$ | Intensity of statoliths/PINs coupling |
| $\mathcal{P}$ | $\mathcal{P} = \frac{W k_{\text{off},1}}{k_{\text{on},1}}$ | Ratio of endocytosis to exocytosis (pool number) |

## 3 RESULTS

### 3.1 Steady gravitropic response: revisiting the sine-law

We first study the gravitropic response predicted by our model in the case of a steady inclination of the plant *θ*, for long timescales when the system reaches a steady state. This situation corresponds to the usual protocol for measuring the sensitivity of plant to gravity under steady condition, when the plant is suddenly inclined to a fixed angle *θ* and its curvature (or tip angle) measured over time. After a transient, the rate of change of curvature is found to be constant (Chauvet et al., 2016), which enables to measure the steady gravitropic response 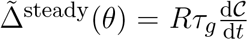 (see Material and Methods) for each imposed angle *θ*. For many plants, this relationship between the gravitropic response and the inclination angle has sine-like shape (the response is null for *θ* = 0° or *θ* = 180° and maximal for *θ* = 90°) and is called the ‘sine-law’ in the literature (Sachs, 1887; Larsen, 1969; Iino et al., 1996; Galland, 2002; Dumais, 2013). Below, we determine the steady gravitropic response 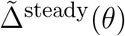 predicted by the model (eqs. 1, 5, 7) and compare with measurements of the sine-law obtained previously for wheat coleoptiles over a wide range of angles (Chauvet et al., 2016).

In the steady regime, the auxin transport equation (eq. 5) reduces to: d*J/*d*x* = 0, where 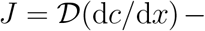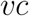 is the auxin flux. For impermeable boundaries at *x* = 0 and *x* = 2*R*, the flux is null and the auxin concentration profile is given by:

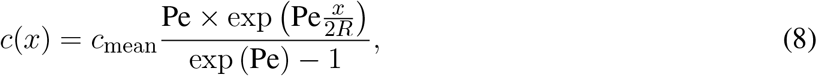

where 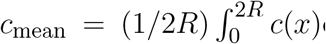 is the mean concentration of auxin and Pe is the Peclet number comparing advection to diffusion and given by:

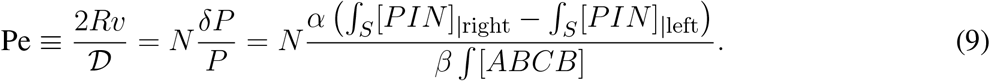

For Pe ≪ 1, the profile is linear and the auxin level in the middle of the stem is unchanged, whereas for Pe ≫ 1 the profile is strongly asymmetric with most auxin concentrated on the right, corresponding to the lower side of the shoot (Fig. 5). From this steady profile of auxin, the gravitropic response can be computed as 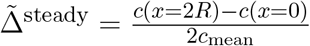 (eq. 1), which gives:

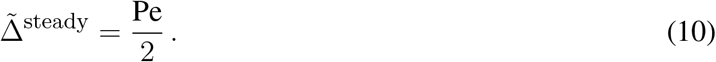

**Figure 5.**
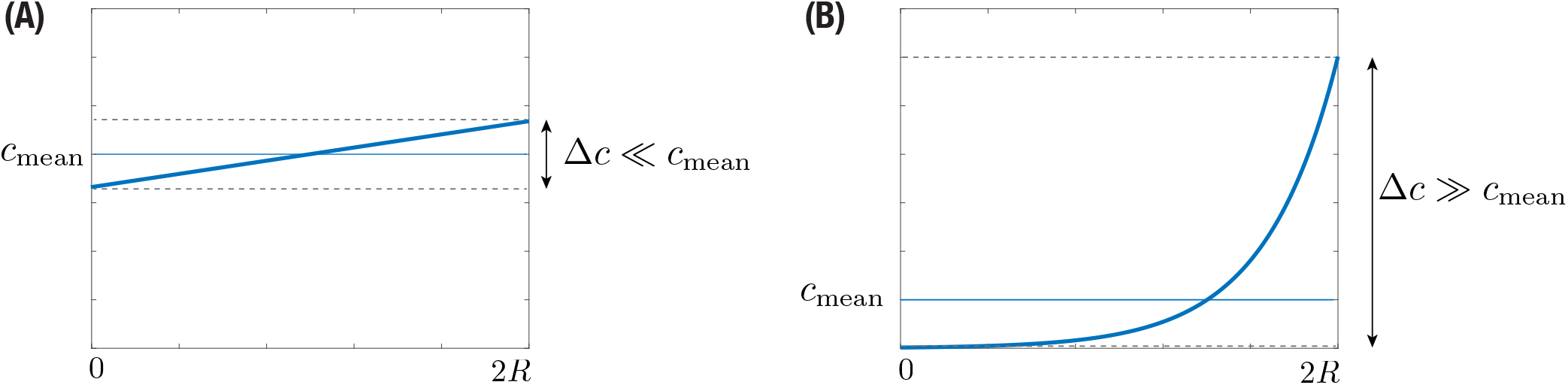
Stationary auxin profile for **(A)** small and **(B)** large Peclet number.

In the steady state, the gravitropic response of the plant is thus given by the value of the Peclet number. Previous measurements in various plant species representative of land angiosperm showed that 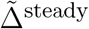 is typically of the order 1 (Chauvet et al., 2016) (for e.g. in wheat coleoptile, the maximal value of 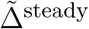 obtained for a 90 degrees inclination is about 0.7), meaning that the Peclet number is typically of the order 1. Therefore, the auxin profile across the shoot is expected to be close to linear (Fig. 5A) (and not a pronounced exponential, Fig. 5B). A consequence is that growth, which was assumed proportional to the auxin concentration, varies also linearly from one side of the shoot to the other during the gravitropic response.

The next step to determine 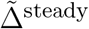 is to compute the Peclet number given by equation (9), i.e., the relative distribution of PIN between the left and right side of the cell. In the steady regime, the concentration of PIN attached to a membrane not covered by statoliths 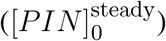, or covered by statoliths 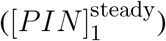, is given by (see eq. 7):

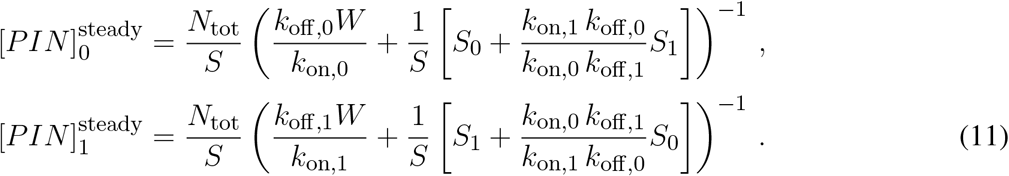

Noting 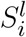 (resp. 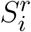) the surface of the left (resp. right) side of the cell not covered (*i* = 0) or covered (*i* = 1) by statoliths, we have 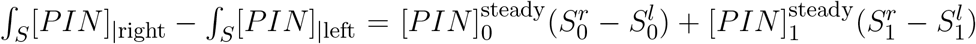. Finally, since 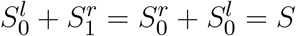 where *S* is the lateral surface of the cells and using equation (9), we have:

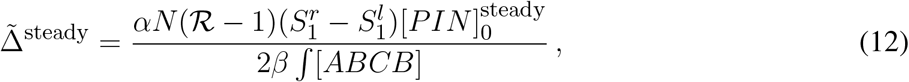

with

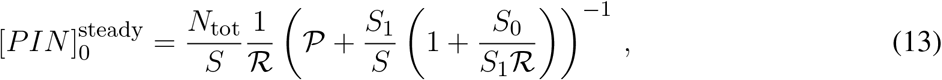

and:

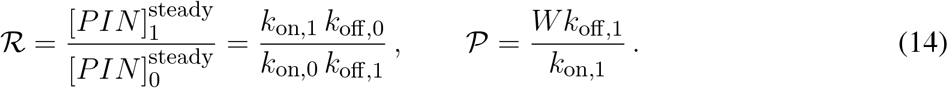

The expressions (12–14) show that the steady gravitropic response is proportional to 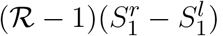. For *θ* > 0 as in figure 1, the difference 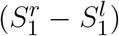 is positive since statoliths sediment toward the right side of the cell. Therefore, to obtain a ‘normal’ gravitropic response (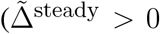, i.e., a larger auxin concentration at the bottom side of the shoot), the ratio 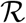 must be larger than 1. The parameter 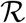 is the ratio between the concentration of PINs in a zone with statolith and in a zone without statolith, hence the larger 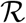, the more PINs in the region with statoliths compared to region without statolith (see Fig 4C). The other main parameter controlling the gravitropic response is the pool number 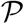 characterizing the ratio of endocytosis to exocytosis, where 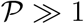 corresponds to the infinite-pool regime and 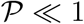 corresponds to the limiting-pool regime (see Fig 4B).

Once 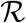 and 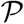 are set, the final step is to compute how the different surfaces covered (and not covered) by the statoliths vary as function of the inclination angle *θ*. To this end, one has to know the final position of statoliths when a cell is tilted. Recently, we addressed this question and showed that statoliths at the bottom of statocytes behave like an effective liquid on long timescale, due to the agitation of statoliths by the cell activity (Bérut et al., 2018). Therefore, the final free surface of the statoliths pile is horizontal, as sketched in figure 3. This key feature of the flowing behavior of statoliths allows us to reduce the computation of the surfaces touched or not by the statoliths (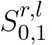, *S*_0_ and *S*_1_ in eqs. 12, 13) to a purely geometrical problem, which depends on three parameters: the angle of inclination *θ*, the aspect ratio of the cell *H/W* and the initial aspect ratio of the statolith pile *H*_0_*/W* (see Fig. 3). The corresponding relationships are given in Appendix B.

Figure 6 presents the typical steady gravitropic response 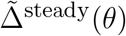 predicted by the model (eqs. 12, 14, 14) as a function of *θ*, in the case of an infinite pool 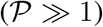 or a limiting pool 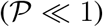, and for two extreme values of 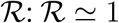 (red curve, low influence of statoliths on PIN binding) and 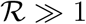 (blue curve, strong influence of statoliths on PIN binding). The geometry used for the cell aspect ratio and the statoliths pile ratio is taken from experimental observations of wheat coleoptile statocytes, with a typical aspect ratio *H/W* = 2.5 and *H*_0_*/W* = 1/2.5. In the infinite pool regime (Fig. 6A), the gravitropic response presents a convex shape with a strong peak close to 90° whatever the value of 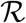, in disagreement with the usual sine-law shape. The amplitude of this peak in this regime also strongly depends on the cell geometry: the more elongated the cell, the higher the peak because in this case more lateral surface is available for PIN attachment without any PIN dilution (see Fig 4B). By contrast, in the limiting-pool regime (Fig. 6B,C), the response strongly depends on 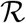, the peak at 90° being much smaller for 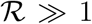 than for 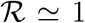. Interestingly, the response in this case also depends on whether PIN can attach on every sides of the cell (apical/basal/lateral binding) or only on the lateral sides (lateral binding case), as attachment to the ‘useless’ apical and basal sides contribute to deplete PIN from the available pool. Overall, we see that only one case is compatible with the concave ‘sine-law’ shape observed experimentally: a limiting pool of PIN with lateral binding and 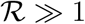 (blue curve in Fig. 6C).

**Figure 6.**
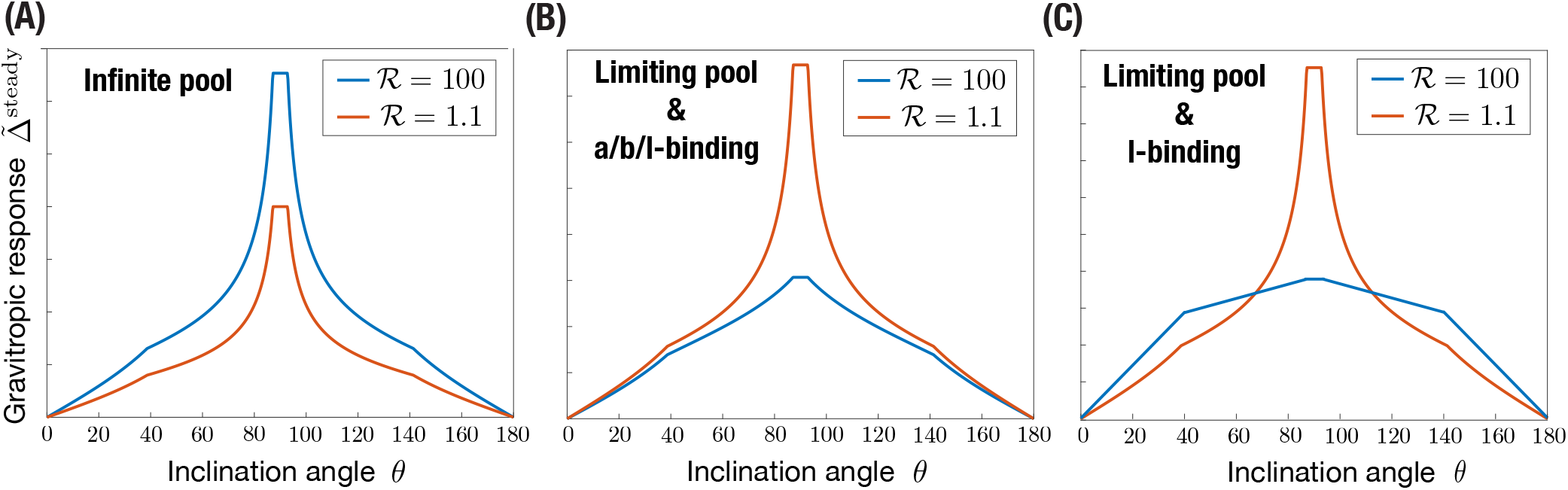
Steady gravitropic response 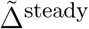 (arbitrary amplitude) as a function of the inclination angle *θ*, for 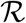 either large or close to 1, in the case of **(A)** infinite pool 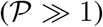, **(B)** limiting pool 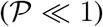 with apical/basal/lateral binding, **(C)** limiting pool 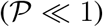 with lateral binding only. Note that in the case of an infinite pool, results are the same in the a/b/l-binding or l-binding case. Geometrical parameters used are *H*_0_ = 4*d*, *W* = 10*d*, *H* = 25*d* where *d* stands for the diameter of a statolith.

In the following, we thus assume that PIN recycling occurs in the limited-pool regime 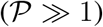, with lateral binding only. In this limit, and using expression of the surfaces given in Appendix B, the steady gravitropic response given by equations (12, 13, 14) can be written as:

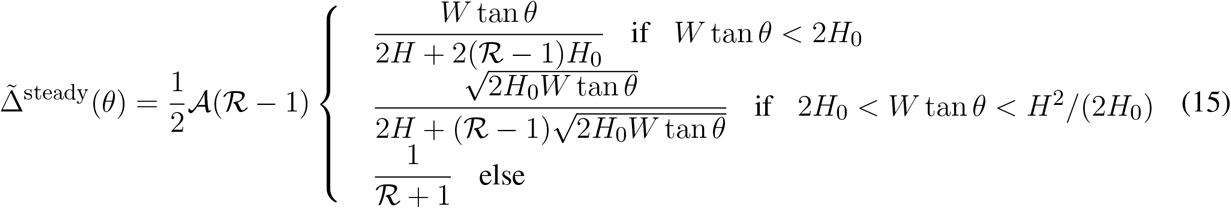

with 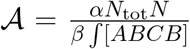. The different conditions stand for cases where statoliths are totally absent of the left side, or totally covering the right side (see Fig. 7). Once the geometry of the cell and of the statoliths pile are fixed, the predicted gravitropic response depends on two dimensionless parameters: 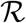 and 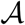. Figure 7 presents the experimental measurements of the ‘sine-law’ obtained by Chauvet et al. (2016) on wheat coleoptiles, together with the best fit of the data using a least-square method. Reasonable agreement between theory and experiments is obtained with 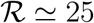 (or larger values as the shape converges in this case) and 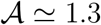 1.3. It is interesting to note that within our position-sensor framework, the predicted steady gravitropic response is not a simple ‘sine-law’, but rather a piece-wise curve with an overall concave shape. This law is also not universal and can be affected by several anatomical and physiological properties, such as the geometry of the cell *H/W*, the amount of statoliths *H*_0_*/W*, or the molecular signaling machinery (embedded in the parameters 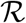 and 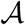).

**Figure 7.**
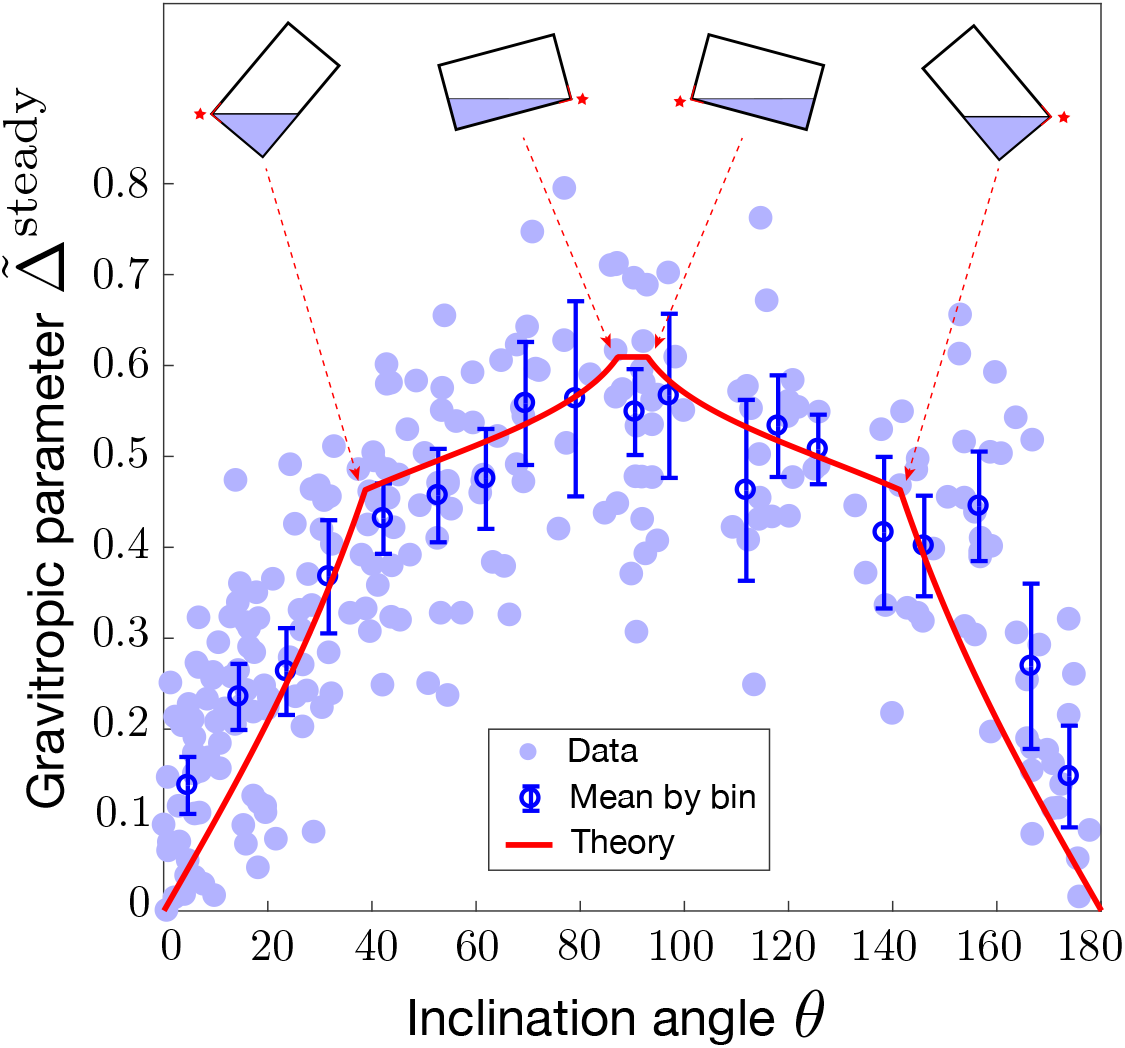
Modified sine-law 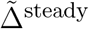 as a function of the inclination angle *θ*. Comparison between the model prediction for 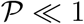 (limiting pool) and l-binding (eq. 15 with 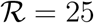 and 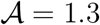, red line) and experiments on wheat coleoptiles (symbols, Chauvet et al. (2016)). Geometrical parameters used in the model for the statocyte are *H*_0_ = 4*d*, *W* = 10*d*, *H* = 25*d* where *d* stands for the diameter of a statolith. Error bars are the mean value of the data by binning the [0,180] interval into 20 boxes.

### 3.2 Transient gravitropic response: dose-response law

The previous results deal with the steady gravitropic response obtained when the gravity stimulus (the angle of inclination *θ* of the plant) is permanent. We now turn to the study of the transient gravitropic response, i.e., when the system has not yet reached the steady state. This situation typically corresponds to ‘dose-response’ like experiments, in which the gravity stimulus is applied during a transient time Δ*T* only. Using such protocol on wheat coleoptiles, Chauvet et al. (2019) revealed the existence of an intrinsic ‘memory’ time *τ*_memory_ in the gravitropic response. For Δ*T* ≫ *τ*_memory_, the response was constant and equal to the steady response 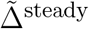. However, for Δ*T* ≲ *τ*_memory_, the response was smaller and became proportional to Δ*T* (Fig. 8). The memory time *τ*_memory_ ∼ 15 min identified in these experiments was longer than the sediment time of statoliths (~2 min) but shorter than the growth timescale (hours). It thus reflects a temporal process in the gravitropic signaling pathway that remains to be identified. We address below this question in the framework of the model.

**Figure 8.**
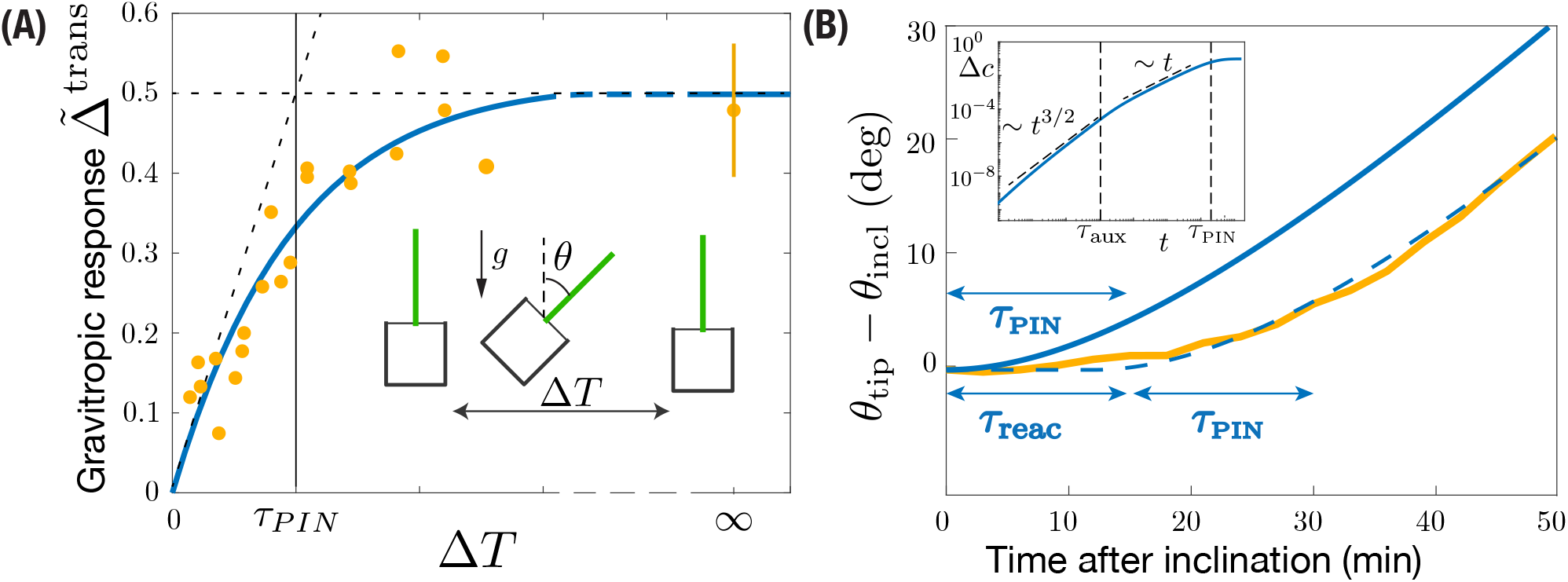
Gravitropic response to a transient inclination (dose-response like protocol). **(A)** Maximal gravitropic response reached during the dynamics as function of the inclination time Δ*T* for *θ* = 45°. The blue solid line is the model prediction using *τ*_PIN_ = 13 min (*τ*_aux_*/τ*_PIN_ and *τ*_1_*/τ*_PIN_ ≪ 1, other parameters are fixed as in Fig. 7). Symbols correspond to the aggregation results of (Chauvet et al., 2019) under 1*g* and 3*g*. **(B)** Evolution of the tip angle after an inclination *θ* = 50° predicted by the model with the same parameters (blue solid line) and in the experiments of Chauvet et al. (2016) (orange thick line). The predicted model must be shifted by a constant time *τ*_reac_ = 13 min to match the experimental curve (dashed blue line). Inset: early time behavior of the gravitropic response predicted by the model in log-log scale.

The set of equations (5)–(7) give a complete description of the transient gravitropic response (we assume that sedimentation of statoliths is fast enough that the surfaces in equation 7 can be computed from their equilibrium values – see Appendix B). Two different typical times control the dynamics: *τ*_aux_, describing the transport of auxin across the tissue of length 2*R*, and *τ*_PIN_, describing the dynamics of PIN at molecular scale. The time scale associated to auxin transport *τ*_aux_ can be estimated using equation (5) for a constant coefficient of diffusion 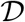 and transport velocity *v* (i.e., PIN distribution). In this case, relaxation towards the stationary profile (eq. 8) is exponential and occurs on a time scale set by the inverse of the shortest non-vanishing eigen-mode of equation (5) (Mohsen and Baluch, 1983):

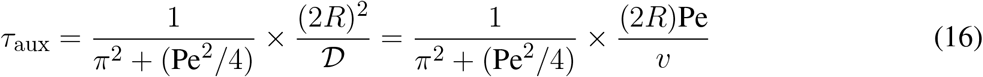

Estimating the Peclet number from the steady gravitropic response (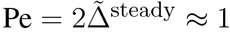, see Fig. 7), and the auxin transport speed *v* from measurements of the speed of auxin pulses in plant tissues (*v* ≃ 3 *μ*m/s, Goldsmith (1977); Rashotte et al. (2003)), gives *τ*_aux_ ≃ 30 s (taking 2*R* ~ 1 mm). This is much shorter than the memory time *τ*_memory_ ~ 15 min evidenced in dose-response experiments, and even shorter than the statoliths sedimentation time. This suggests that the dynamics of the gravitropic response is controlled by *τ*_PIN_, rather than by the auxin diffusion time *τ*_aux_.

The time scale *τ*_PIN_ is set by the slowest characteristic time of the system (eq. 7) describing PIN dynamics. From the eigenvalues of this linear system, two timescales are obtained, which are solutions of:

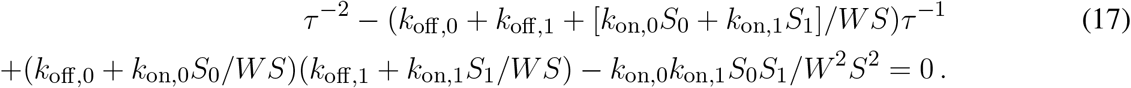

In the limiting pool case (*WSk*_off,i_*/k*_on,i_*S*_*i*_ ≪ 1), the two solutions are:

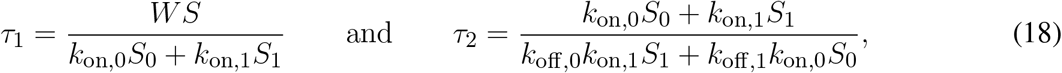

with *τ*_1_ ≪ *τ*_2_. Therefore, the slowest timescale of the gravitropic signaling pathway, which sets *τ*_memory_, should be given by *τ*_PIN_ = *τ*_2_. Note that when preferential attachment in region with statoliths is achieved via a strongly increased attachment rate (*k*_on,1_ ≫ *k*_on,0_), and not by change of detachment rate (*k*_off,0_ ≃ *k*_off,1_), then 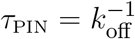. Conversely, if it is achieved by decreased detachment rate (*k*_off,0_ ≫ *k*_off,1_) and not by change of attachment rate (*k*_on,0_ ≃ *k*_on,1_), then 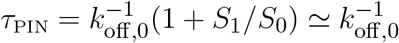. Remarkably, in this last case, the equilibration time of PIN is not controlled by the slowest rate of detachment (*k*_off,1_), but by the fastest one (*k*_off,0_), due to the limiting pool.

To check these predictions, we solve the model for a transient gravitropic stimulus that reproduces the protocol of Chauvet et al. (2019). The inclination *θ* is set to 45° for a transient time Δ*T* and then put back to zero, the gravitropic response being defined as the maximal auxin gradient reached during the dynamics: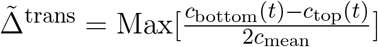. Figure 8A compares the experimental data of Chauvet et al. (2019) with the prediction of the model for *τ*_aux_*/τ*_PIN_ ≪ 1, using the parameters 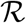 and 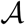 already fixed by the steady response (see Fig. 7). Good agreement is obtained using *τ*_PIN_ = 13 min as the only fitting parameter. This result shows that *τ*_PIN_ is playing the role of the memory time evidenced by the experiments of Chauvet et al. (2019). If Δ*T* ≪_PIN_, PIN transporters have not enough time to rearrange before the end of the stimulus, and the response is weak. Conversely if Δ*T* ≫ _PIN_, PINs have time to rearrange and reach their steady repartition before the end of the stimulus, and the response is maximal, similar to the one of a permanent stimulus.

We finish our analysis by investigating the full temporal dynamics of the gravitropic response after a sudden inclination *θ* = 50°. Figure 8B presents the time evolution of the shoot curvature 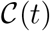 (or similarly *θ*_*tip*_(*t*) for small curvatures) predicted by the model for *τ*_PIN_ = 13 min (blue curve), together with the experimental data of Chauvet et al. (2016) (yellow curve). The time scale of the curvature change *τ*_PIN_ is well captured by the model. However, to match the experiments, the model has to be shifted in time by a constant time *τ*_reac_ ≈ 13 min. Such delay or reaction time of the gravitropic response after the stimulus was already noticed by Chauvet et al. (2019), but does not seem to be captured by the model. Actually, a careful analysis of the temporal behavior of the model (assuming *τ*_1_ ≪ *τ*_aux_ ≪ *τ*_PIN_ = *τ*_2_) shows that the auxin gradient at early times increases as Δ*c* ~ *t*^3/2^ (so that 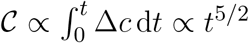) as long as *t* < *τ*_aux_, before varying as Δ*c* ~ *t* for *τ*_aux_ < *t* < *τ*_PIN_ (Inset of Fig. 8B). These scaling laws are thus not compatible with a very flat initial response.

## 4 DISCUSSION

In this paper, we have derived a multi-scale model of plant gravitropism which links the different steps of the gravitropic signaling pathway: (i) the initial intracellular perception of gravity by statoliths, (ii) the transduction of this physical signal into a biochemical signal through the reorganization of PINs at the membrane of statocytes, (iii) the intercellular signal transmission via auxin transport and (iv) asymmetric organ growth. The main originality of the model lies in the mechanistic link we propose between the statoliths position and the dynamics of PINs, based on the recent position-sensor hypothesis (Pouliquen et al., 2017). This basic assumption coupled to auxin transport and growth in an idealized tissue made of a one-dimensional array of cells recovers several major features of the gravitropic response of plants.

### 4.1 A new interpretation of the sine-law of plant gravitropism

The first main result concerns the steady gravitropic response to a permanent gravity stimulus, 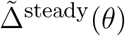. For many plants, this response takes the form of an inclination-dependent law with a sine-like shape, called for this reason the sine-law (Sachs, 1887; Larsen, 1969; Iino et al., 1996; Galland, 2002; Dumais, 2013). This sine-law has long been interpreted in terms of a force sensor mechanism, for the projected weight of the statoliths on the lateral surface of the cell varies with the sine of the inclination angle (Audus, 1969; Barlow, 1995). However, recent experiments showing that the response is independent of the gravity intensity have dismissed this force-sensing hypothesis, calling for a new interpretation of the sine law (Chauvet et al., 2016; Pouliquen et al., 2017). A key result of the model is that it predicts an inclination-dependent steady gravitropic response 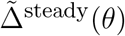 without invoking a force-based mechanism. In the model, the initial gravitropic stimulus is the statoliths position at the cell membrane, not their weight. Since statoliths behave on long time like a liquid (Bérut et al., 2018), their position in steady state is a purely geometrical cue, which depends only on the cell inclination. As a result, the steady response predicted by the model depends on the inclination but not on the gravity intensity, in agreement with the observations.

In the model, the actual shape of the gravitropic response 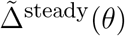 is never a pure sine law. It depends on several parameters, related either to geometric factors, such as the aspect ratios of the statocytes and of the sedimented statoliths pile, or to molecular processes: intensity of coupling between statoliths position and PINs through parameter 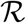; ratio of endocytosis to exocytosis through parameter 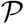; ratio between the conductance of PIN carriers to the total conductance of auxin transporters through parameter 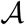. For elongated cells and shallow statoliths piles such as those of wheat coleoptile statocytes, the shape of the response tends to be sine-like only in the case of a strong coupling between the statoliths and PINs 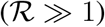, and for a number of PINs conserved along the cell membrane (limiting pool regime, 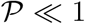). This latter assumption is common in models of auxin transport (Runions et al., 2014; Retzer et al., 2019), while the strong coupling assumption is compatible with the large asymmetry in PINs localization observed upon gravity stimulation (Friml et al., 2002; Harrison and Masson, 2008; Kleine-Vehn et al., 2010). Interestingly, although gravity is absent from the model, the gravitropic response depends on the amount of statoliths through the geometrical aspect ratio of the statoliths pile *H*_0_*/W*. The model could thus account for previous experiments using starch-less and starch-excess mutants, which showed a modified gravitropic response when the number of statoliths is changed (Kiss et al., 1997; Vitha et al., 2007; Pouliquen et al., 2017). Finally, it is worth noting that the model assumes that the statoliths form a static pile at the bottom of the cell, while statoliths actually exhibit a dynamic and random agitation due to the interaction with the cytoskeleton (Sack et al., 1986; Saito et al., 2005; Nakamura et al., 2011; Bérut et al., 2018). We might expect this agitation to reduce the averaged contact time between the statoliths and the cell membrane, thereby decreasing the coupling between statoliths and PINs. It would be interesting to extend the model in order to incorporate such effect of agitation on the gravitropic response. The model could then be compared with the behavior of agravitropic mutants in *Arabidopsis thaliana* like *sgr9*, whose weaker response is likely associated to an abnormally strong agitation of the statoliths (Nakamura et al., 2011).

Overall, our model suggests that the classical sine-law of plant gravitropism might not be universal, as its shape and amplitude could depend on several anatomical and physiological parameters. Full measurements of the gravitropic response of plants over a wide range of inclination are scarce and mostly performed on shoot coleoptiles. It would be interesting to perform systematic measurements of the sine-law on other plant organs (root, stem), to see if the shape of the sine-law is affected by different statocyte geometries and tissue organization.

### 4.2 The gravity-independent memory process in dose-response laws is likely associated to PIN dynamics

The second main result of our study concerns the gravitropic response to a transient gravity stimulus. For a sudden inclination applied at time *t* = 0, the model predicts that the response reaches the steady response 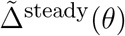 only after a time large compared to a ‘memory’ timescale *τ*_memory_, corresponding to the slowest timescale introduced in the model. Therefore, when a stimulus is applied only during a transient time Δ*T* < *τ*_memory_, a weaker gravitropic response is predicted, following a dose-response like law. In the model, the memory time is not associated with the sediment time of the statoliths, which is assumed to be much shorter than the other timescales of the gravitropic signaling pathway (a valid assumption for a gravity intensity like Earth gravity). Our model is therefore compatible with the recent experiments of Chauvet et al. (2019) performed on wheat coleoptiles, which show a dose-like behavior of the gravitropic response with a memory time *τ*_memory_ independent of gravity. The model also provides the explicit origin of this memory process, which was postulated in Chauvet et al. (2019). In the model, two different processes can lead to the temporal filtration of the initial signal associated with statoliths position: auxin transport across the tissue and the dynamics of PIN turnover at the molecular scale. Our study suggests that the limiting process is actually controlled by PIN dynamics (*τ*_memory_ = *τ*_PIN_), auxin transport being too fast to account for the memory time measured experimentally (~ 10-20 min). Interestingly, visualization of the PIN3 auxin efflux carrier in root columella cells after a sudden change in the gravity vector indicates a time scale of about 10 min for complete relocation (Friml et al., 2002), a duration very close to the memory time measured by (Chauvet et al., 2019). Although our model successfully captures the existence and origin of a gravity-independent memory process in the signaling pathway, it is not able to describe the delay time *τ*_reaction_ ~ 10 min observed between the application of the stimulus and the first gravitropic response (Chauvet et al., 2019). This delay may have different origins: the time needed to reorganize the cytoskeleton implied in the transport of PIN carriers, the time needed by a PIN to go from the pool towards the plasma membrane, or the time of incorporation of a PIN into the membrane. It is worth noting that a similar timescale of about 10 min was identified in the gravity-sensing columella cells for the internalization of PIN3 from the plasma membrane into vesicles (Kleine-Vehn et al., 2010). Further experiments combining a transient stimulus with pharmacological and genetic approaches would be needed to confirm the key role of the PIN turnover timescale in the gravitropic response. Besides, it would be interesting to see what is the influence of this hierarchy of time-scales on the response of a plant to oscillatory stimulus (e. g. wind) or on the dynamic competition between gravitropism and proprioception (Bastien et al., 2013).

### 4.3 Back to the statoliths/PIN coupling assumption

We conclude by discussing the possible origin of the coupling between statoliths and PINs, which is at the core of our model. Although the respective roles of statoliths and PIN auxin transporters in plant gravitropism are well established, the link between the two is still not clear (see Nakamura et al. (2019) for a recent review of the possible molecular actors involved). In our model, we have used a very general hypothesis for this coupling based on the recent finding that the relevant gravitropic stimulus is the statoliths position inside the gravisensing cells (Chauvet et al., 2016; Pouliquen et al., 2017; Bérut et al., 2018). We have postulated that statoliths in contact with the cell membrane bias the exocytosis and endocytosis rate of PIN recycling, therefore inducing an asymmetry of PIN distribution when statoliths reposition in response to plant inclination. Our results suggest that this interaction between PINs and statoliths is strong, as large values of the parameter 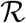 are needed to match the experimental gravitropic response. This interaction between statoliths and PINs could involve a complex molecular pathway that remains to be unveiled (Strohm et al., 2014). However, more direct mechanisms of interaction could occur. For example, statoliths could modify PIN vesicle-mediated transportation to the membrane by modifying the architecture of the actin cytoskeleton. Another possibility would be that internalization of PINs from the membrane is reduced by the presence of statoliths, for example by simple steric effects. Indeed, direct visualizations reveal a length scale of ~ 1 *μ*m for the endosome formation, which is not far from statolith size (Kleine-Vehn et al., 2010). Interestingly, such direct interaction of statoliths and the cytoskeleton machinery was put forward as an explanation of gravitropism in rhizoids and protonemata, such as the single-cell alga *Chara* (Sievers et al., 1996). A last possibility could be that PINs cluster in regions where there is no statolith, while they do not cluster when statoliths are present. Indeed, for a conserved number of PIN proteins, clustering reduces the efflux efficiency, as this diffusion process scales not linearly but as the square-root of the number of carriers (Bénichou and Voituriez, 2014; Valet et al., 2019). Such a clustering has been highlighted by Kleine-Vehn et al. (2011), but, to our knowledge, no comparison has been done between regions of the membrane in contact or not in contact with statoliths. Further studies would be needed to discriminate between these molecular mechanisms.

## AUTHOR CONTRIBUTIONS

N.L., O.P., and Y.F. designed research. N.L. built and analyzed the model with inputs from O.P. and Y.F. N.L., O.P., and Y.F. wrote the paper..

## ACKNOWLEDGMENTS

The authors would like to thank Valérie Legué and Bruno Moulia for the many stimulating discussions we have continually had on this subject.

## APPENDIX A CASE OF STATOLITHS ONLY PRESENT ON A RING

Let be a central region between *l* and 2*R* − *l* with no statolith (and constant efflux rate *P*_0_). We can redo the calculation leading to eq. (4) in this geometry. Clearly, equations will be similar in each region, with *v* = 0 in the central one. We have to take care to the continuity of the flux at the jonction of each regions. We can show that *J* (*l*) = (*P* +*δP*)*c*(*l*^−^)−*P*_0_*c*(*l*^+^) and *J* (2*R*−*l*) = *P*_0_*c*((2*R*−*l*)^−^)−*Pc*((2*R*−*l*)^+^). At steady state, we thus get a jump of auxin concentration: *c*(*l*^+^) = (*P* +*δP*)*/P*_0_ *c*(*l*^−^) and *c*((2*R*−*l*)^+^) = *P*_0_*/P c*((2*R*−*l*)^−^). In the limit of small Peclet number, we thus get 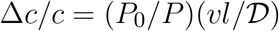.

## APPENDIX B EXPRESSION OF THE GEOMETRIC PARAMETERS AS A FUNCTION OF *θ*

We give here the value of the surfaces 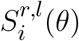 that are necessary to compute the Peclet number as a function of the angle (see eq. 15). As statoliths are solid, the shaded area in Fig. 3 is conserved and is thus equal to *H*_0_*W*. We note *θ*_1_ = arctan 2*H*_0_*/W* the angle for which the horizontal level meets the lower left corner, and *θ*_2_ = arctan *H*^2^/(2*H*_0_*W*) the one for which it meets the upper right one (we assume *H*_0_ < *H/*2). Elementary geometry calculations give:

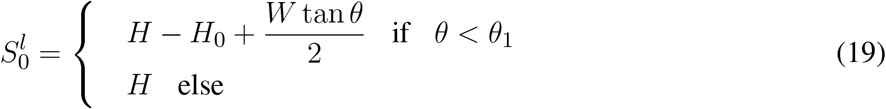

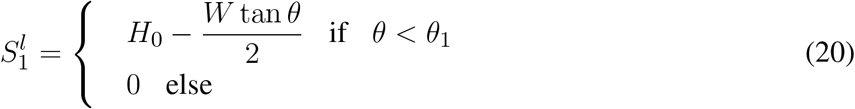

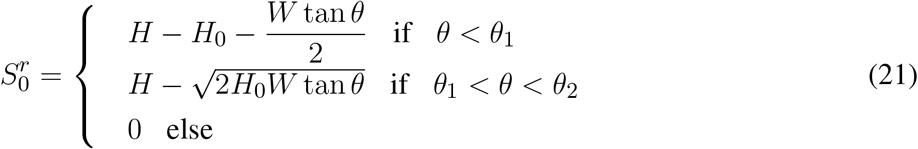

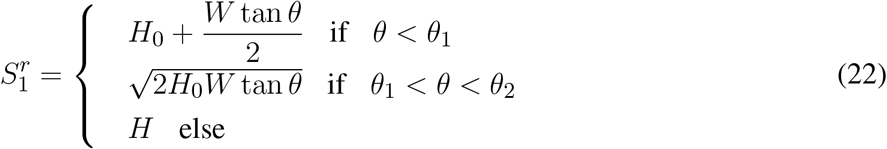

In the case of apical binding, this leads to

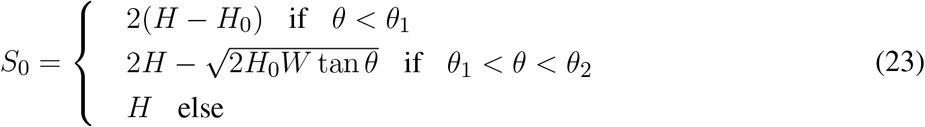

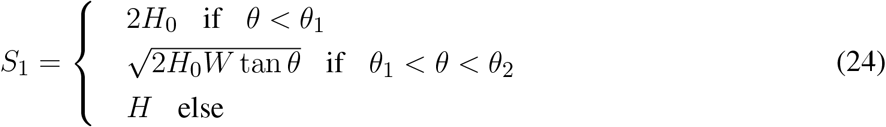

In the case of apical/basal/lateral binding, the expressions for the total surfaces are slightly modified due to attachment at apical and basal side.

